# CREATION AND CHARACTERIZATION OF A RECOMBINANT MAMMALIAN ORTHOREOVIRUS EXPRESSING HUMAN EPIDERMAL GROWTH FACTOR RECEPTOR 2 PEPTIDES

**DOI:** 10.1101/2023.06.12.544655

**Authors:** Nicole A. Jandick, Cathy L. Miller

**Author notes:** Corresponding author, Address: VMRI #3, 1907 ISU C. Drive, Ames, IA, 50011.

## Abstract

Mammalian orthoreovirus (MRV) is an oncolytic virus that has been tested in over 30 clinical trials. Increased clinical success has been achieved when MRV is used in combination with other onco-immunotherapies. This has led the field to explore the creation of recombinant MRVs which incorporate immunotherapeutic sequences into the virus genome. This work focuses on creation and characterization of a recombinant MRV, S1/HER2nhd, which expresses three human epidermal growth factor receptor 2 (HER2) peptides (E75, AE36, and GP2) known to induce HER2 specific CD8+ and CD4+ T cells. We show S1/HER2nhd expresses HER2 peptides in infected cells and on the virion, and infects, replicates, and reduces HER2+ breast cancer cell survival. The oncolytic properties of MRV combined with HER2 peptide expression holds potential as a vaccine to prevent recurrences of HER2 expressing cancers.

## Introduction

Genetically modified oncolytic viruses are increasingly being approved for human use against several cancer types. Oncorine (H101), a serotype 5 adenovirus with genetic deletions in the E1B-55k and E3 genes, was approved in China in 2005 (Liang, 2018). Talimogene laherparepvec (T-VEC), a modified herpes simplex virus type 1 (HSV-1) with deletions in infected cell protein 34.5 (ICP34.5) and ICP47, as well as insertion of the gene expressing granulocyte macrophage colony-stimulating factor (GM-CSF), was approved in the USA in 2015 (Andtbacka et al., 2015; Andtbacka et al., 2019; Kaufman et al., 2017). Additional genetically engineered viruses that encode exogenous sequences, such as Pexa-Vac, a thymidine kinase-deleted poxvirus that is loaded with the GM-CSF gene (Parato et al., 2012) and Prostvac-VF, a vaccinia and fowlpox prime-boost vector combination that delivers a prostate cancer specific antigen (PSA) and three immunostimulatory factors, are under investigation (Heo et al., 2013). In each of these virus gene delivery systems, the virus vector must be mutated such that it is less virulent than the wildtype (WT) virus. Some oncolytic virus systems, such as mammalian orthoreovirus (MRV), are clinically benign in their WT state, and may be useful for the production of genetically engineered virus systems to deliver exogenous genetic material to the tumor environment.

MRV was discovered to replicate to high levels in transformed cells relative to non-transformed cells over 40 years ago (Duncan et al., 1978). Further investigation of MRV exposed tumor regression in RAS activated cells after MRV infection (Coffey et al., 1998; Strong et al., 1998). Since this discovery the virus has been investigated pre-clinically and in clinical trials as both a monotherapy and as part of a dual therapy regimen. Clinical trials have shown limited success using MRV as a monotherapy, therefore researchers have turned their focus to utilizing MRV in combination with other therapeutics (Chaurasiya et al., 2021). With the recent advent of a plasmid-based MRV reverse genetics (RG) system (Kobayashi et al., 2007; Kobayashi et al., 2010), an additional approach gaining favor is the design of recombinant viruses that express proteins and peptides that enhance the oncolytic and anti-tumor properties of MRV. The complexity of packaging a segmented genome and the MRV requirement for the expression of one protein from every gene segment has made recovering recombinant viruses challenging. Nonetheless, progress is being made and several groups have been able to recover stable recombinant MRVs that express exogenous sequences.

Successful recombinant MRV viruses engineered to express exogenous sequence can be separated into two general categories, one in which two proteins are expressed from a single mRNA, and one in which a single fusion protein is created by cloning genes expressing MRV proteins in frame with non-viral sequences. In the first category, 2A sequences, which are a naturally occurring stretch of amino-acids ending in Proline/Glycine found in a number of viruses (Luke et al., 2008), are added to MRV gene segments to drive the production of 2 proteins from one MRV mRNA. During protein translation the 2A sequence triggers the ribosome to skip due to the formation of the glycyl-prolyl peptide bond (Donnelly et al., 2001). This results in the release of the first protein, and ribosome reinitiation. Use of the 2A sequence results in the addition of a number of amino acids (aa) on the upstream protein, and a proline on the downstream protein (Donnelly et al., 2001). Using this method with porcine teschovirus-1 2A (P2A), a near-infrared fluorescent protein iRFP720, for the potential use of tumor imaging, was added to the M1 gene of MRV (Ogawa et al., 2022). However, by passage 3, the gene encoding the infrared fluorescent protein was lost (Ogawa et al., 2022). Another group added sequence encoding the NanoLuc protein to the MRV L1 gene to allow live imaging of tumor cells (Kanai et al., 2019). While the resulting recombinant virus was shown to replicate in cells, and NanoLuc was highly expressed in mice, passage stability was not discussed (Kanai et al., 2019). In another study a modified σ1 protein, junction adhesion molecule A (JAM-A) independent-3 (*jin-3)* (van den Wollenberg et al., 2012), containing a single mutation in σ1 at aa 196 that increases MRV entry via sialic acid, was used to generate multiple constructs using the 2A approach. Using the *jin-3* σ1 expressing construct, iLov, a fluorescent protein (van den Wollenberg et al., 2015), early region 4 open reading frame 4 (E4orf4), an adenoviral protein known to induce apoptosis (Kemp et al., 2018), and human and mouse GM-CSF (Kemp et al., 2019) were incorporated in place of the head region of σ1. Recombinant viruses using these modified constructs were rescued and found to exhibit variable stability upon long term passage.

The second approach, in which non-viral protein coding sequences are fused in frame with MRV protein sequence, is limited to proteins that retain functionality when non-viral sequences are added. Thus far, viral proteins μNS and σ1 have been modified using this approach. Viral non-structural protein, μNS, forms viral factories where transcription, translation, and virion assembly occur during viral replication (Dales, 1963; Desmet et al., 2014; Kniert et al., 2022; Miller et al., 2010). Viral protein σ1, is the attachment protein of MRV and is the most varied between serotypes (Nibert et al., 1990). σ1 can bind both JAM-A and sialic acid residues for entry into the cell and is thought to first bind onto sialic acid residues and be pulled closer to the cell membrane, allowing attachment to JAM-A (Barton et al., 2001a; Barton et al., 2001b). Our lab added a tetracysteine-tag, which is 6 aa in length, to the non-structural μNS coding sequence at the carboxyl (C)-terminus and at aa 545 and were able to recover recombinant virus (Bussiere et al., 2017). The C-terminal tagged virus accumulated a second-site mutation that did not influence functionality of μNS or the tetracysteine tag, but when the tetracystine tag was added to μNS at aa 545, the protein lost functionality and was not stable at the sequence level over long term passage (Bussiere et al., 2017). Another stable recombinant MRV was create by adding a 9 aa arginylglycylaspartic acid (RGD) motif, which binds integrins on the surface of cells, onto σ1 in order to increase binding to cells that do not express JAM-A (Kawagishi et al., 2020). The RGD motif was added in frame at different sites throughout the σ1 protein and resultant recombinant MRVs were found to be stable, replicate to the same level as WT and were infectious in JAM-A knock-out cancer cells (Kawagishi et al., 2020). This work showed additional sequence could be added into the loops of the head domain of σ1, however when more than one RGD sequence was added, viral replication was reduced compared to WT and furthermore, deletion mutants occurred over time suggesting the introduced sequence was not stably retained in the virus. With these factors in mind, we set out to identify small immunogenetic sequences that could be added to MRV.

MRV has been investigated as a potential therapeutic in a multitude of tumor environments and has been shown to replicate well in tumor cells that express epidermal growth factor receptor (EGFR/HER1) and was even thought at one time to use EGFR/HER1 for entry (Strong and Lee, 1996). A closely related growth factor receptor and an EGFR/HER1 binding partner is HER2. HER2 overexpression is seen in a variety of cancers, including breast, bladder, gastric, ovarian, salivary gland, endometrial, pancreatic, and non-small-lung cancer (Iqbal, 2014). HER2 overexpressing cancers have rapid cell growth and increased rate of relapse (Slamon et al., 1987). Several peptides (E75, GP2, and AE36) derived from the HER2 protein have been explored as potential therapeutic vaccines against HER2+ breast cancer (BC) (Brown et al., 2020; Clifton et al., 2016; Mittendorf et al., 2012; Mittendorf et al., 2014; Sears et al., 2011). E75 is found in the extracellular domain, GP2 in the transmembrane domain, and AE36 in the intracellular domain of HER2 (Fisk et al., 1995; Peoples et al., 1995). Each of these peptides have been shown in clinical trials to induce CD8+ T cells that target HER2 expressing cells, and AE36 has also been shown to induce a CD4+ T cell response (Brown et al., 2020). Thus far, the peptides have been tested in combination with GM-CSF to induce the immune response, however some studies have shown lower T cell responses and induction of inhibitory myeloid-derived suppressor cells (MDSCs) with GM-CSF use (Serafini et al., 2004; Slingluff et al., 2009). In this study, we created a recombinant MRV expressing GP2, AE36, and E75 peptides fused in frame with the tail domain of σ1 as a potential therapeutic vaccine against HER2 overexpressing cancers and characterized the recombinant MRV in HER2 overexpressing BC cell lines. Our data suggest that this recombinant MRV binds to and replicates in cells to levels similar to a gene-matched control MRV, is stable over 3-6 passages, and expresses HER2 peptides in infected cells and on the surface of the virion.

## Materials and Methods

### Cells, viruses, antibodies, and reagents

AU-565 (estrogen receptor (ER)-, progesterone receptor (PR)-, HER2 3+, Ki-67 95%), BT-474 (ER+, PR+, HER2 3+, Ki-67 70%), and ZR-75-1 (ER+, PR+/-, HER2 2+, Ki-67 80%) (Subik et al., 2010) cell lines were maintained in Roswell Park Memorial Institute (RMPI) 1640 medium (Gibco) supplemented with 10% fetal bovine serum (Atlanta Biologics) and penicillin (100 I.U./ml) streptomycin (100 μg/ml) solution (Mediatech). The MCF7 (ER+, PR +/-, HER2 0, Ki-67 90%) (Subik et al., 2010) cell line was maintained in Minimum Essential Media (Gibco) supplemented with 10% fetal bovine serum (Atlanta Biologics), 1% human insulin (Sigma-Aldrich), and penicillin (100 I.U./ml) streptomycin (100 μg/ml) solution (Mediatech). L929 cells were maintained in Joklik modified minimum essential medium (Sigma-Aldrich) supplemented with 2% fetal bovine serum (Atlanta Biologics), 2% bovine calf serum (HyClone), 2 mM L-glutamine (Mediatech), and penicillin (100 I.U./ml) streptomycin (100 μg/ml) solution (Mediatech). BHK-T7 cells were maintained in Dulbecco’s Modified Eagle Medium (DMEM) (Gibco) supplemented with 10% fetal bovine serum (Atlanta Biologics), 1X MEM non-essential amino acids (Cytiva) and penicillin (100 I.U/ml) streptomycin (100 μg/ml) solution (Mediatech). Our laboratory stock of Type 3 Dearing Cashdollar (T3D) was obtained from the laboratory of M.L. Nibert and originated in the laboratory of L.W. Cashdollar (Parker et al., 2002). Primary antibodies used were as follows: polyclonal rabbit α-μNS as described in (Qin et al., 2011), polyclonal rabbit α-Flag (Cell Signaling DYKDDDK Tag Antibody; #2368), monoclonal mouse α-Flag (Sigma-Aldrich 41106000 M2-Flag), α-T3D virion antisera, and α-T1L virion antisera (Virgin et al., 1988). Secondary antibodies used were Alexa 488- and 594-conjugated donkey α-mouse or α-rabbit IgG antibodies (Invitrogen Life Technologies; #A-21202, #A-21207), and goat α-mouse or α-rabbit IgG alkaline phosphatase (AP)-conjugate antibodies (Bio-Rad Laboratories, #1706520, #1706518).

### Plasmid construction and RG assays

pT7-S1/HER2nhd (no head domain) was constructed using pBacT7S1-T3D (Addgene, (Kobayashi et al., 2007) as a backbone plasmid. A double-stranded DNA (gBlock, Integrated DNA Technologies) containing WT Type 3 Dearing (T3D) S1 sequence until nucleotide (nt) 767 (aa 252) with a nt substitution from 596 to 598 of GGG to CGA (aa G196R) followed by the coding sequence for HER2 peptides GP2, E75, and AE36, a Flag tag, and a stop codon was designed and purchased. Both pBacT7S1-T3D and the gBlock were digested using NcoI and SbfI for 2 h at 37°C, isolated using DNA Clean & Concentrator-5 (Zymo Research) then ligated using T4 DNA Ligase (New England BioLabs Inc.). The resultant plasmid contained the gBlock inserted before the last 124 nucleotides of WT S1 which includes the ‘C’ Box (Roner and Roehr, 2006) that is thought to be necessary for S1 gene segment packaging. After transformation, bacterial colonies were isolated, amplified, extracted using Plasmid Miniprep Kit (Zippy Research Corporation) and digested with SacI at 37°C for 4 hours to confirm insertion of the gBlock, before being isolated using DNA Clean & Concentrator-5 (Zymo Research) and sent for sequencing. Sequencing was confirmed using 5’-GCTATTGGTCGGATGGATCCTCGCC −3’ as the forward primer, and 5’-TGAAATGCCCCAGTGCCG −3’ as the reverse primer, allowing coverage of the full sequence of the pT7-S1/HER2nhd plasmid. Once sequence was confirmed, the plasmid was retransformed, amplified, and extracted using Plasmid Maxi Kit (QIAGEN), according to manufacturer’s protocol.

MRV RG assays were used to generate S1/HER2nhd virus. In addition to the pT7-S1/HER2nhd, pT7-L1-M2T1L, pT7L2-M3T1L, pT7-L3-S3T1L, pT7-M1T1L, pT7-S2T1L, and pT7-S4T1L (Addgene) described in (Kobayashi et al., 2007; Kobayashi et al., 2010) were transfected into BHK-T7 cells at 0.4μg/μl. Accessory plasmids pCAG-D1R and pCAG-D12L (Addgene) as described in (Kanai et al., 2017) were transfected at 0.2μg/μl, and pCAG-FAST-p10 (Addgene) as described in (Kanai et al., 2017) was transfected at 0.005μg/μl. TransIT-LT1 (Mirus Bio) was used in a 3:1 ratio of TransIT-LT1 to μg of total plasmid. Transfected cells were incubated for 5 days at 37°C. 5 days post transfection (d p.t.) cells were subjected to three freeze/thaw cycles before being collected in glass 1-dram vials. Following the same RG assay protocol, a gene matched control virus containing all T1L gene segments except S1, which was WT T3D (LS1D), was generated using pBacT7-S1T3D (Addgene) instead of pT7-S1/HER2nhd. To generate recombinant T1L, the same RG assay protocol was used, using pBacT7-S1T1L (Addgene) for the S1 segment.

### Plaque assay, virus passage, and virus purification

L929 cells were plated at 1.2 x 10^6^ per well in 6 well plates. 24 hours post plating (h p.p.), serial dilutions of RG assay lysates or replication assay lysates in phosphate-buffered saline (PBS) (137 mM NaCl, 3 mM KCl, 8 mM Na_2_HPO_4_, 1.5 mM KH_2_PO_4_, pH 7.4) with 2 mM (PBS-MgCl_2_) MgCl_2_ (PBS-MgCl_2_) were added to L929 cells. Incubation lasted 1 hour at room temperature, with gentle shaking every 15 minutes, to keep cells hydrated. After 1 hour, L929 monolayers were overlaid with 2 ml of m199 (Irvine Scientific) containing 2 mM L-glutamine (Mediatech), and penicillin (100 I.U./ml) streptomycin (100 μg/ml) solution (Mediatech), 0.225% NaHCO_3_, 2 mg/ml trypsin and 1% Bacto-Agar (BD, 214010) and incubated for 3 days at 37°C and 5% CO_2_. 3 days post infection, plaques were counted and collected from RG assays as passage (P)0 viral stocks. P0 were subjected to three freeze/thaw cycles before being added to T25 flasks of BHK-T7 monolayers and incubated at 37°C until 80% cell death to create P1 stocks. P1 T25 flasks were subjected to three freeze/thaw cycles before 500 μl P1 was added onto T75 flasks of BHK-T7 monolayers and incubated at 37°C until 80% cell death to create P2 stocks. This was repeated to obtain P3s-P10s for stability assays. As passage number increased, 100 μl-200 μl of previous passage was used to generate the next passage. For purification, P3s were added to BHK-T7 cell monolayers in three-layer flasks. P4s were incubated for three days before being collected and purified as previously described using Vertrel XF (DuPont) instead of Freon (Furlong et al., 1988; Mendez et al., 2000).

### RNA extraction and sequencing

1.88 x 10^11^ viral particles of purified S1/HER2nhd or LS1D as determined by absorbance (A260) = 2.1 × 10^12^ particles, (Smith et al., 1969), or 1 ml of unpurified passages of S1/HER2nhd or LS1D were subjected to RNA extraction via Trizol-LS (Life Technologies) according to manufacturer’s instructions. Briefly, virus samples were homogenized with Trizol-LS. Chloroform was added and the RNA phase was separated and precipitated then washed with isopropanol and ethanol. After RNA extraction, samples were either collected in 2x protein loading buffer (125mM Tris-HCl[pH 6.8], 200mM dithiothreitol [DTT], 4% sodium dodecyl sulfate [SDS], 0.2% bromophenol blue, 20% glycerol) to be separated on 8% SDS-polyacrylamide gel electrophoresis (PAGE) or subjected to reverse transcriptase-polymerase chain reaction (RT-PCR) using Superscript IV (Invitrogen Life Technologies), to create cDNA and then DNA to amplify for sequencing.

### Infection, binding assay, and replication assay

For replication assays AU-565, BT-474, MCF7, and ZR-75-1 were infected with S1/HER2nhd or LS1D at an MOI of 0.1 for 1 hour at 37°C with shaking every 15 minutes. S1/HER2nhd and LS1D were diluted in PBS-MgCl_2_ so equal volume was added onto cells during the 1-hour incubation. Multiplicity of infection (MOI) was determined by first measuring cell infectious units (CIU) on each cell line, as described in (Qin et al., 2011). Once CIU/ml of the viral preparation was determined, the amount of virus to add to cells was calculated by multiplying the number of cells by the desired MOI, which gives the total CIU needed to infect at that MOI. 0, 24, 48, 72, and 96 hours post infection (h p.i.) samples were subjected to three freeze/thaw cycles to stop infection. Plaque assays were performed to determine viral titer at each timepoint. For binding assays, L929 cells were incubated at 4°C for 30 minutes before being mock infected or infected with 5 x 10^10^ viral particles as determined by absorbance at (A260) =2.1 x 10^12^ particles, (Smith et al., 1969) or infected with an MOI of 1 of S1/HER2nhd or LS1D, diluted in PBS MgCl_2_. One hour after incubation at 4°C, supernatants were collected on ice, cells were washed 3 times with PBS before being collected in 1 ml of PBS-MgCl_2_. Samples were stored at 4°C until plaque assays were performed to determine viral titer. For all other infections, AU-565, BT-474, MCF7, and ZR-75-1 were infected with S1/HER2nhd or LS1D at an MOI of 1.

### Immunoblotting and Coomassie Blue staining

2x protein loading buffer was added to 6 x 10^10^ CIU of S1/HER2nhd or LS1D at a 1:1 ratio. Samples were heated to 95°C for 10 minutes before being loaded onto 10% SDS-PAGE gels. For immunoblot analysis, viral proteins separated by SDS-PAGE at 100V for 3 hours were transferred to nitrocellulose by electroblotting at 100V for 40 minutes, then blocked in 5% milk in Tris-buffered saline (20mM Tris, 137mM NaCl [pH 7.6]) with 0.1% Tween-20 (TBST) for 1 hour. Nitrocellulose membranes were then incubated for 18 hours with primary antibodies in 5% milk in TBST, and secondary antibodies in 5% milk in TBST for 2 hours. Membranes were washed with 1x TBST 3 times and incubated for 5 minutes 3 times between antibody incubations and before addition of NovaLume Atto Chemiluminescent Substrate AP (Novus Biologicals). Membranes were then imaged on a ChemiDoc SRS Imaging System (Bio-Rad Laboratories) and Quantity One imaging software (Bio-Rad Laboratories) was used to capture images. For Coomassie Blue analysis, viral proteins separated by SDS-PAGE were stained with 0.025% Coomassie Brilliant Blue R-250 (Bio-Rad Laboratories), 30% isopropanol, 20% methanol for 1 hour, and destained with water for 24-48 hours.

### RNA electrophoresis

2x protein loading buffer was added to 5 x 10^10^ viral particles of T1L, T3D, S1/HER2nhd, or LS1D at a 1:1 ratio. Samples were heated at 65°C for 10 minutes before being loaded onto an 8% SDS-PAGE gel. The gel was run at 16 mA for 12 hours, before being incubated for 1 h in water with 3X GelRed (Phenix Research Products) and imaged on ChemiDoc XRS imaging system (Bio-Rad Laboratories) and Quantity One imaging software (Bio-Rad Laboratories).

### Immunofluorescence

AU-565, BT-474, MCF7, or ZR-75-1 cells were plated at 3 x 10^5^ on glass coverslips. The next day, cells were mock-infected or infected with LS1D or S1/HER2nhd at MOI=1. 24 h p.i. cells were fixed with 4% formaldehyde for 20 minutes. Samples were washed with PBS 3X before being permeabilized with 0.2% Triton X-100 in PBS for 5 minutes. Samples were washed with PBS 3 times, blocked with 1% bovine serum albumin in PBS (PBSA) and 10 minutes later washed 3 times with PBS. Primary antibodies were diluted in PBSA and added to the samples before being incubated with rocking for 1 hour. Samples were washed 3 times with PBS before being incubated with rocking with secondary antibodies diluted in PBSA for 1 hour. After 3x PBS washes, cover slips were fixed onto glass slides using Prolong Gold antifade reagent with DAPI (4,6-diamidino-2-phenylindole dihydrochloride; Invitrogen Life Technologies). Coverslips were examined on Zeiss Axiovert 200 inverted microscope equipped with fluorescence optics. Representative images were taken on Zeiss AxioCam MR color camera using AxioVision software (4.8.2) and prepared using Adobe Photoshop and Illustrator software (Adobe Systems).

### Viability assay

AU-565, MCF7, and ZR-75-1 were plated at 1 x 10^3^ and BT-474 at 5 x 10^3^ in 96 well plates. Cells were mock infected or infected with S1/HER2nhd or LS1D at MOI=1, in triplicate and incubated at 37°C for 24, 72, or 120 hours, at which time Cell Counting Kit (CCK)-8 reagent (APExBIO, K1018) was added to the cells according to manufactures instructions. Two hours post-addition of CCK-8 reagent, samples were analyzed at 450nm on Glomax Multi Detection Plate Reader (Promega).

### Statistical analysis

For replication assay graphs, error bars were generated by calculating standard deviation using =stdev() function in Excel. All other graphs show mean, as larger line, and one standard deviation on either side of mean, as the smaller lines, both calculated using JMP. P-values were calculated using a 2 sided student’s T test, assuming equal variance using =ttest() function in Excel.

## Results

### Design and construction of S1/HER2nhd

To create a recoverable and stable recombinant MRV expressing immunogenic HER2 peptides, we took advantage of prior findings in which the σ1 head domain of the *jin-3* S1 mutant was replaced with other coding sequences (Kemp et al., 2018; Kemp et al., 2019; van den Wollenberg et al., 2012). We created a plasmid, termed pT7-S1/HER2nhd, encoding T3D S1 nts 1-767 (up to aa 252) with substitution from nts 596-598 of GGG to CGA (aa G196R) followed by the coding sequence for HER2 peptides GP2, E75 and AE36, a Flag tag, a stop codon and finally the ‘C’ box, which consists of the 3’ terminal 124 nts thought to be needed for packaging of S1 (Roner and Roehr, 2006) (Fig 1A). S1/HER2nhd is 557 nts shorter than WT S1, and σ1/HER2nhd is predicted to be 33 kDa compared to WT σ1 at 50.6 kDa.

**Fig 1.**
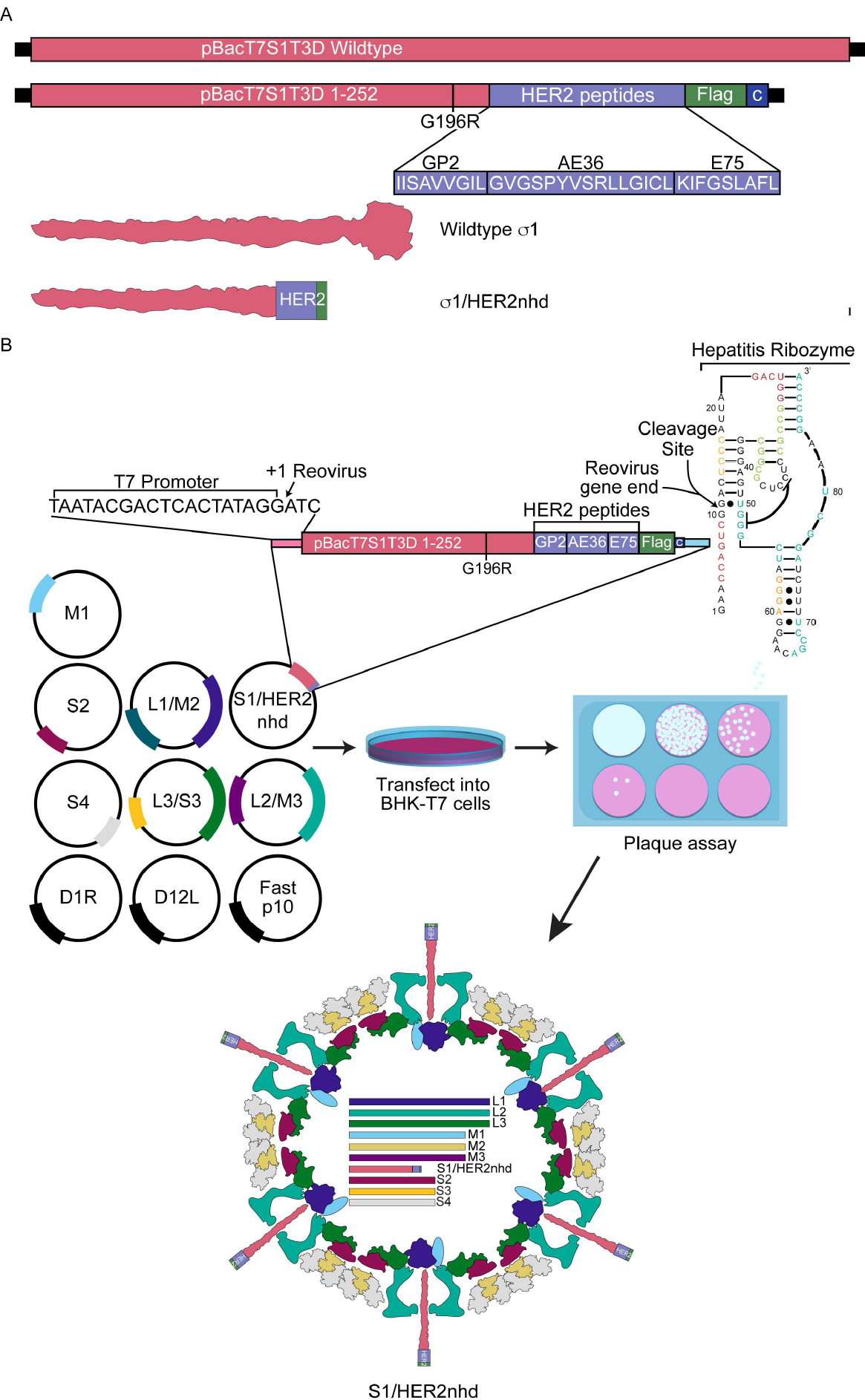
Design and generation of S1/HER2nhd. A) Comparison of the WT pBacT7S1T3D with pBacT7S1/HER2nhd. pBacT7S1/HER2nhd contains WT Type 3 Dearing (T3D) S1 sequence until nt 767 (aa 252) with a nt substitution from 596 to 598 of GGG to CGA (aa G196R) followed by the coding sequence for HER2 peptides GP2, E75 and AE36, a Flag tag, a stop codon, and the C box, comprised of the last 124 nucleotides of S1 needed for S1 packaging. Beneath the plasmid construct illustration are the GP2, E75, and AE36 peptide sequences, and a schematic of the structure for σ1 and σ1/HER2nhd proteins. B) Reverse Genetics protocol used to generate S1/HER2nhd virus. WT T1L MRV plasmids pT7S2, pT7S4, pT7M1, pT7L1/M2, pT7L2/M3, and pT7L3/S3, along with pBacT7T3DS1/HER2nhd, and accessory plasmids D1R, D12L and Fast p10 were transfected into BHK-T7 cells and incubated at 37°C for 5 days. Samples were subjected to three freeze/thaw cycles and plaque assays were performed. Recombinant virus plaques were collected and passaged and RT-PCR products from viral RNA were sequenced to confirm S1/HER2nhd.

pT7S1-HER2nhd, or pBacT7-S1T3D, were transfected in place of pT7-S1T1L with plasmids encoding the 9 other WT T1L MRV gene segments and accessory plasmids in the MRV RG assay (Fig 1B). A pT7-S1T1L positive control was also included in the assay. Five days post transfection, cells were subjected to three freeze/thaw cycles, before being serial diluted and analyzed via plaque assays, in which we found that we were able to generate recombinant virus plaques in pT7-S1T1L, pBacT7-S1T3D and pT7-S1/HER2nhd-containing RG samples. S1/HER2nhd and LS1D generated a similar number of plaques as recombinant S1/T1L from RG assays, however, S1/HER2nhd and LS1D plaques were smaller and more easily detected on day 3 post-plaque assay relative to recombinant S1/T1L plaques. Three plaques were picked from the pT7-S1/HER2nhd samples and passaged until passage 3 for RNA extraction and sequencing to confirm the viruses possessed the expected S1 mutation. Each of the plaques was found to contain the intended mutations at this early passage number, confirming this approach was successful for incorporating S1 gene segments encoding a “headless” σ1 in-frame with a HER2 peptide/Flag tag fusion protein into a replicating MRV.

### S1/HER2nhd virus has expected gene segments and σ1 protein size

To examine the incorporation of the mutant and strain-specific S1 gene segments, purified T1L, T3D, LS1D and S1/HER2nhd virions were separated on an 8% SDS-PAGE gel and the dsRNA genome segments were stained with 3X GelRed. Our expectation was that the S1/HER2nhd L, M, and S2, S3, and S4 segments would migrate with the T1L segments, while the S1/HER2nhd segment, because of the removal of the S1 head domain sequence, would migrate lower on the gel than WT T3D S1. As expected, S1/HER2nhd migrated below S2, S3, and S4, providing proof that the mutated segment was packaged into the virion in its truncated form (Fig 2A). The genotype matched control virus, LS1D showed the expected L, M and S2-S4 segment migration, which matched that of the T1L virus, and the expected S1, which migrated with T3D S1 (Fig 2A). To confirm that the mutated σ1/HER2nhd protein was incorporated into virions, LS1D or S1/HER2nhd virions were separated on SDS-PAGE and proteins were stained with Coomassie. Figure 2B shows that the λ, μ1, σ2, and σ3 proteins for LS1D and S1/HER2nhd viruses all co-migrate, and as expected, the σ1/HER2nhd protein was located at around 33 kDa as a result of the truncation of the σ1 protein. Taken together with the sequencing data, these findings strongly support that the S1/HER2nhd gene segment is incorporated into the virion, and that the σ1/HER2nhd protein is assembled onto the virion, providing proof that our approach in creating this recombinant MRV was successful.

**Fig 2.**
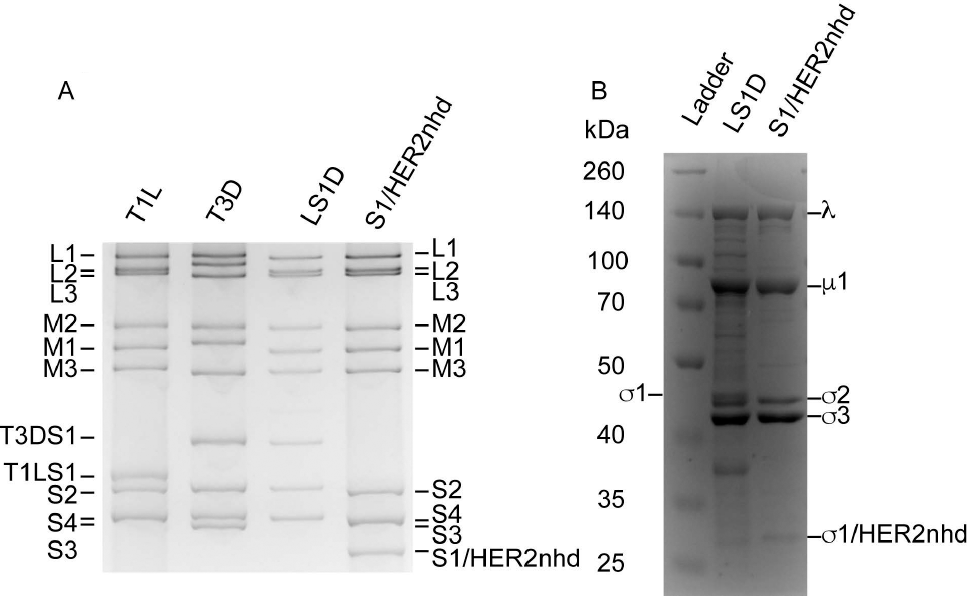
S1/HER2nhd genomic RNA and proteins are of expected size. A) 1.88 x 10^11^ virions of T1L, T3D, LS1D, and S1/HER2nhd were separated on SDS-PAGE and stained with 3X GelRed. 5 x 10^10^ virions of LS1D or S1/HER2nhd were separated on SDS-PAGE and stained with Coomassie Blue dye.

### S1/HER2nhd expresses HER2 peptides on the surface of the virion

One unique aspect of the design of S1/HER2nhd, is that the immunogenic HER2 peptide/Flag fusion is protruding off the virion at the location normally occupied by the head domain of σ1 (Fig 1A, B). The direct presentation of HER2 peptides on the virion may result in B cell activation and antibody production against HER2, in addition to the activation of CD8+ T cells and CD4+ T cells known to be created when these HER2 peptides are expressed with GM-CSF (Brown et al., 2020; Fisk et al., 1995; Holmes et al., 2008). To determine if the σ1/HER2 peptide/Flag fusion resulted in expression of the HER2 peptides on the virion of S1/HER2nhd, purified LS1D and S1/HER2nhd virions were separated on SDS-PAGE, transferred onto nitrocellulose and immunoblotted with antibodies against the Flag tag and the MRV virion. A band between 35 and 25 kDa representative of σ1/HER2nhd was clearly present in the S1/HER2nhd particle sample immunostained with α-Flag antibodies (Fig 3A), which is indicative of expression of the HER2 peptides on the virion of S1/HER2nhd.

**Fig 3.**
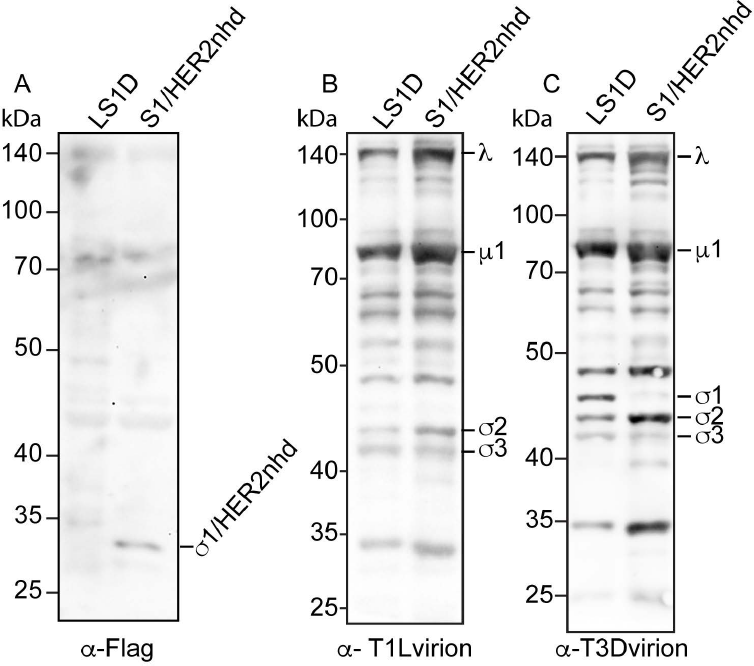
Flag antibodies recognize σ1/HER2nhd on MRV virions. 5 x 10^10^ virions of LS1D or S1/HER2nhd were separated on SDS-PAGE, before being transferred to nitrocellulose and stained with A) rabbit α-Flag polyclonal antibodies, B) rabbit α-TIL virion antisera, or C) rabbit α-T3D virion antisera, followed by HRP-conjugated α-rabbit secondary antibodies, exposure to chemiluminescent reagent, and image capture.

When the nitrocellulose was incubated with antibodies raised against the MRV T1L virion, the expected outer capsid and core proteins for both LS1D and S1/HER2nhd were present (Fig 3B). We additionally probed with antibodies raised against the T3D virion to demonstrate the presence of the T3D σ1 on the LS1D virus (Fig 3C). This antibody also binds the T1L outer capsid and core proteins, likely due to similarities between these proteins in T3D and T1L serotypes (Breun et al., 2001). Altogether, these experiments demonstrate that the σ1/HER2nhd protein containing the HER2 peptides is present on purified virions, strongly suggesting it will be presented to the immune system if used as a therapeutic vaccine against HER2+ BC.

### S1/HER2nhd stability upon passage

Prior research has provided evidence that insertion of exogenous genetic material, or mutation of MRV proteins is not well tolerated by the virus, and that second site mutations and deletions can occur during extensive passaging of recombinant MRV (Bussiere et al., 2017; Kemp et al., 2018; Kemp et al., 2019). To test the stability of the S1/HER2nhd virus, three plaques recovered after RG assay were passaged in BHK-T7 cells 10 times. This process included inoculating T25 flasks of BHK-T7 cells with the P0 of each isolated plaque, growing virus until 80% cell death, subjecting cells to three freeze/thaw cycles, then adding 100μl-500μl, depending on passage number, of each passage cell lysate onto new T75 flasks of BHK-T7 cells. RNA was isolated from each passage cell lysate and subjected to Sanger sequencing using S1-specific primers. As seen in Figure 4A, S1/HER2nhd plaque 1 was stable until passage 6, at which point a stop codon was introduced upstream of the HER2 peptides/Flag tag sequence. This mutation prevents translation of the HER2/Flag fusion, but the RNA sequence encoding HER2/Flag remains in the genome. As passaging was continued, this second-site mutation became the predominant sequence within the virus population. In S1/HER2nhd plaque 2 and plaque 3, stability was rapidly lost with the introduction of a change in the S1/HER2nhd population sequence between passages 3 and 4. These changes again resulted in loss of expression of the HER2/Flag fusion peptide, with plaque 2 containing a partial deletion of the HER2 peptides and Flag tag (Fig 4B) and plaque 3 containing the same stop codon introduced in plaque 1 (Fig 4C). Despite the loss of stability over time we measured in these experiments, we have been able to routinely produce purified S1/HER2nhd virus with no second site mutations in S1 using the standard MRV purification protocol where virus is purified at passage 4.

**Fig 4.**
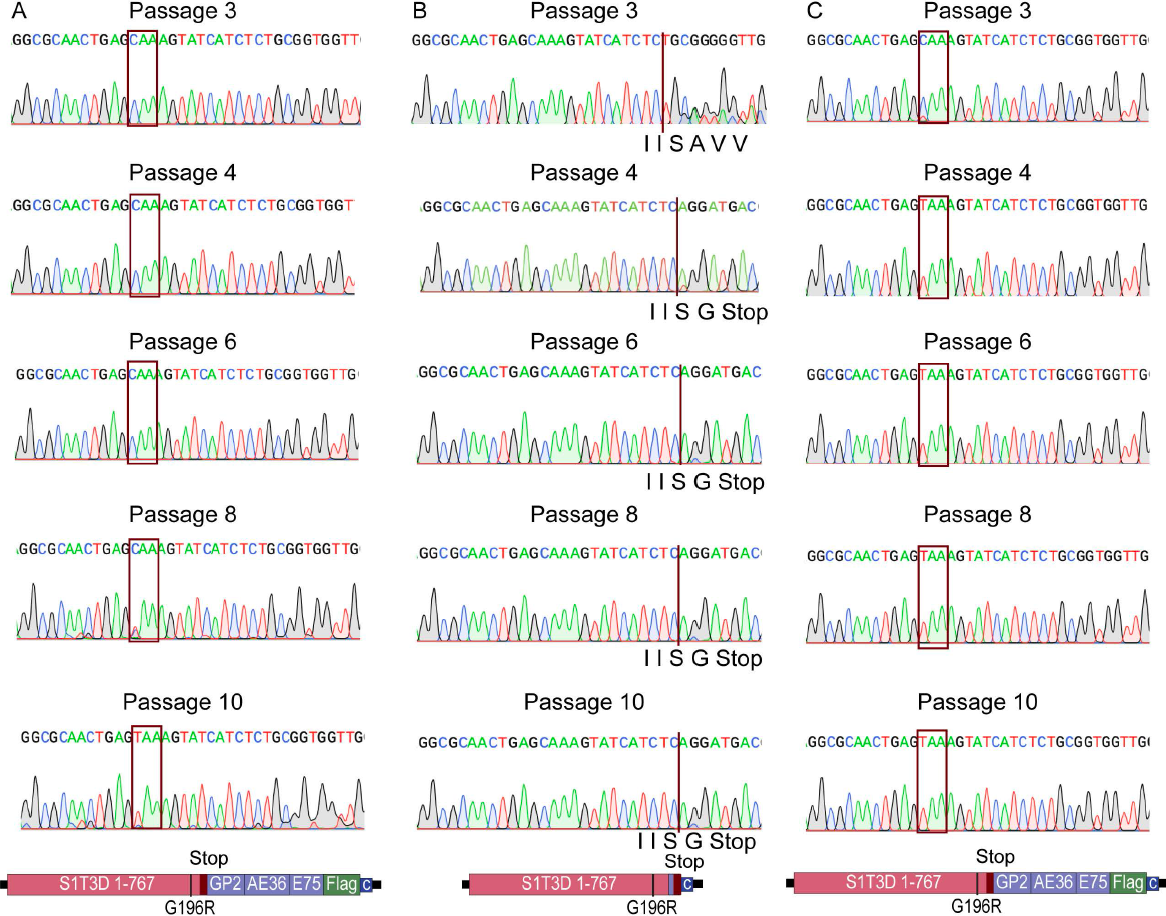
S1/HER2nhd stability upon passage. Three plaques were picked after reverse genetics and passaged 10 times. RT-PCR products from RNA isolated from passage 3, 4, 6, 8 and 10 were sequenced. A) Plaque 1 sequence was stable through passage 6, but at passage 8 the boxed sequence changed from CAA to TAA, introducing a stop codon before the HER2 peptide sequence. B) Plaque 2 contained a partial deletion of the HER2 peptides starting at passage 4, maintaining the “IIS” sequence, but deleting the remaining HER2 peptide/Flag tag sequence and introducing a stop codon. C) Plaque 3 introduced the same stop codon as Plaque 1 in passage 3 which was fully established in the population by passage 4.

### S1/HER2nhd binds L929 cells with same efficiency as LS1D

Replacing the head domain of σ1with the HER2 peptide/Flag fusion will result in loss of binding of MRV to the JAM-A receptor, however, the S1/HER2nhd virus would be expected to continue to bind cells via σ1 attachment to sialic acid residues. However, it is possible that the addition of the HER2 peptide/Flag fusion may interfere with the σ1:sialic acid interaction. To determine the extent by which S1/HER2nhd can bind cells relative to LS1D, L929 cells were mock infected or infected with either 5 x 10^10^ virus particles of LS1D or S1/HER2nhd or at an MOI of 1 of LS1D or S1/HER2nhd at 4°C. After 1 hour, plaque assays were used to analyze virus titer in the supernatant and cell lysate samples. As seen in Fig 5A-C, an average of 8% of input S1/HER2nhd virus was bound onto cells after one hour of incubation, and an average of 6% of input LS1D was bound to cells. There was no significant difference in percent of input virus bound between S1/HER2nhd and LS1D, suggesting that when the same number of particles of S1/HER2nhd or LS1D are exposed to cells for one-hour, similar amounts of virions bind onto the cell surface. When using an MOI of 1, which translates to 1.06 x 10^10^ particles S1/HER2nhd and 1.08 x 10^10^ particles LS1D, we again measured the same percentage of cells with bound virus in S1/HER2nhd and LS1D (Fig 5D-F). These results suggest that the σ1/HER2nhd protein does not have a deficiency in facilitating the binding of virions to L929 cells relative to WT T3D σ1.

**Fig 5.**
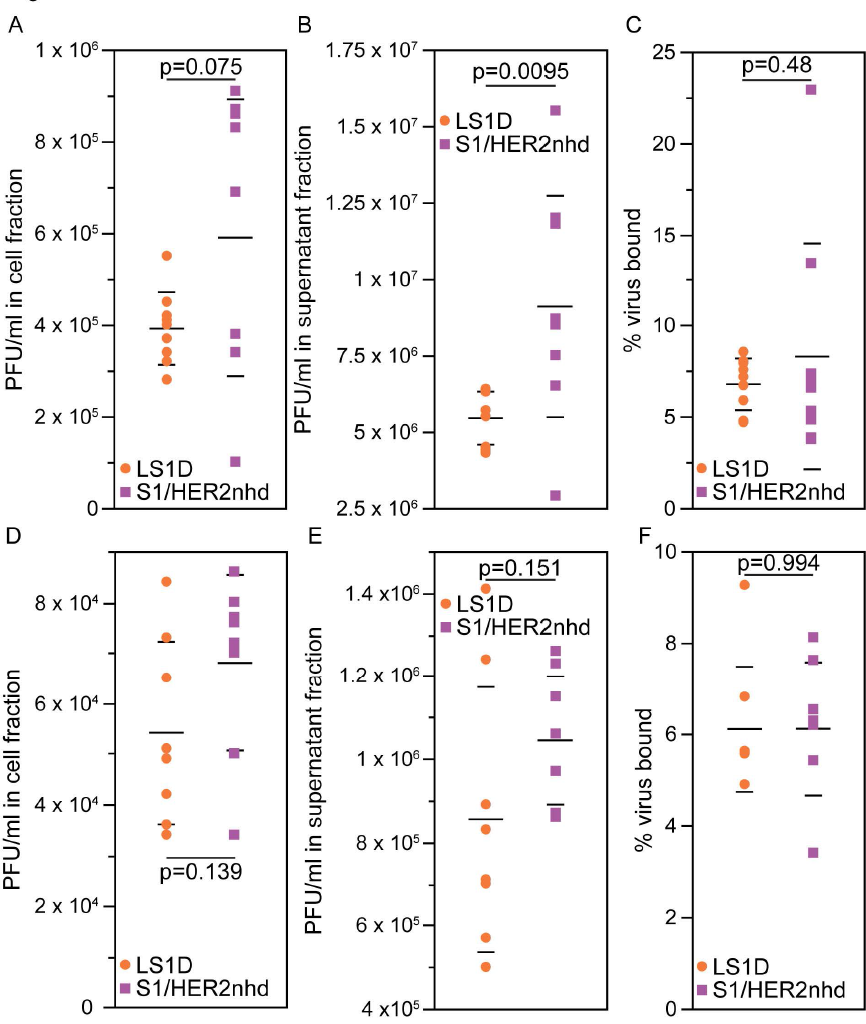
S1/HER2nhd and LS1D bind L929 cells with similar efficiency. L929 cells were mock-infected or infected with A-C) 5 x 10^10^ particles of LS1D or S1/HER2nhd or D-F) MOI=1 LS1D or S1/HER2nhd. Cells were incubated with virus at 4°C for 1 hour, cell and supernatant fractions were collected and analyzed for viral titer using plaque assays. Percent bound virus is also shown (C, F). The means and standard deviation of three replicates are shown and the p-values, calculated using a 2-sided student T test, are as indicated.

### S1/HER2nhd expresses σ1/HER2/Flag fusion in BC cells

In addition to presenting HER2 peptides on the surface of the σ1 molecules on virions, S1/HER2nhd is expected to express HER2 peptides as part of the σ1 fusion protein at high levels inside infected cells, which could then be both presented to the immune system via Class I MHC and released at high levels following lysis of infected cells and subsequently taken up by antigen processing cells. To determine if S1/HER2nhd infects and expresses the σ1/HER2 peptide/Flag fusion protein in BC cells, we infected MCF7, ZR-75-1, AU-565, and BT-474 BC cell lines with LS1D or S1/HER2nhd. At 24 h p.i., immunofluorescence assays were performed using antibodies against MRV nonstructural protein μNS to identify actively infected cells and the Flag tag to measure expression of the σ1/HER2/Flag fusion protein.

In each BC cell line, LS1D and S1/HER2nhd viruses both clearly infected the cells as indicated by μNS expression. However, only S1/HER2nhd infected cells showed cells that were positive for both μNS and for Flag tag immunostaining (Fig 6A-D, lower panels). This finding indicates S1/HER2nhd virus can bind, enter, and express HER2 peptides from the σ1 fusion protein inside BC cells, suggesting that it may be able to activate an immune response against HER2 from within infected cancer cells and following tumor cell lysis.

**Fig 6.**
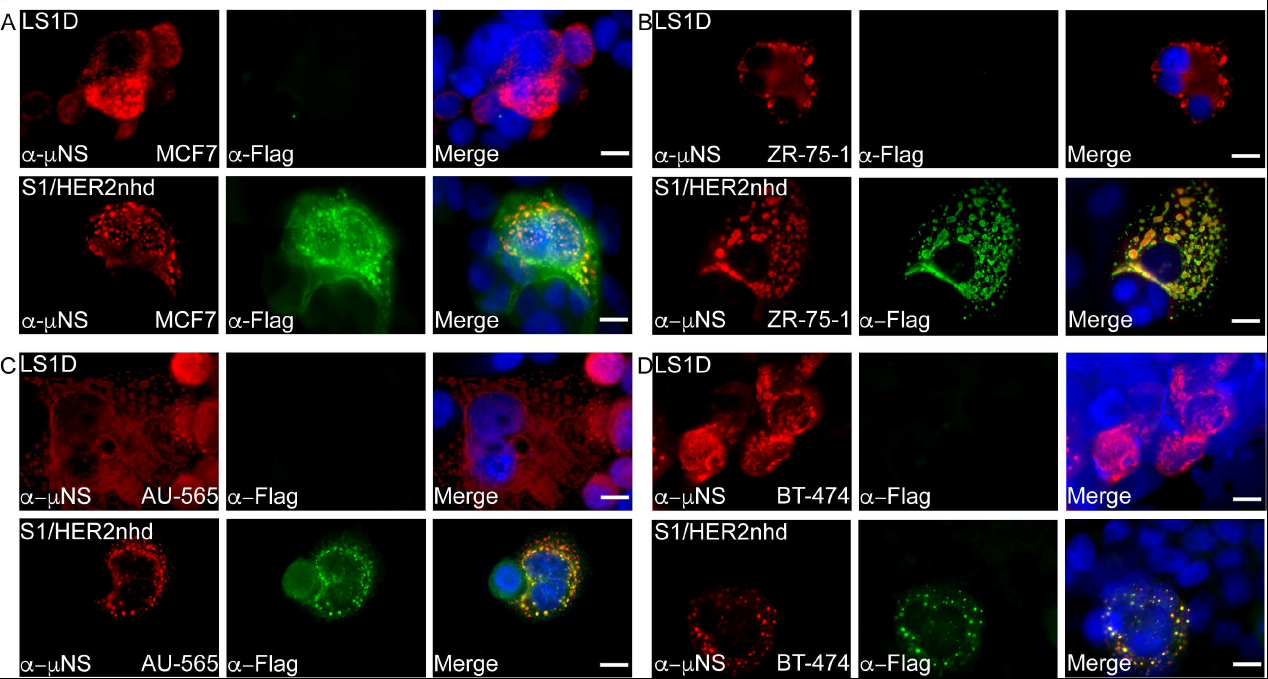
S1/HER2nhd expresses HER2peptide/Flag fusion in infected cells. BC cell lines A) MCF7 (HER2 0), B) ZR-75-1 (HER2 2+), C) AU-565 (HER2 3+) and D) BT-474 (HER2 3+) were infected with LS1D (top) or S1/HER2nhd (bottom) for 24 hours then subjected to immunofluorescent assay, staining with polyclonal α-μNS rabbit (red) and monoclonal α-Flag mouse (green) followed by Alexa 594-conjugated α-rabbit and Alexa 488-conjugated α-mouse secondary antibodies. Cells were counterstained with DAPI (blue) and merged images are shown. Scale Bar =10μm.

### S1/HER2nhd replicates in BC cells

While S1/HER2nhd appears to replicate and form plaques in BHK-T7 and L929 cells (Fig 4, Fig 5) and can bind, enter, and express proteins in BC cells (Fig 6), we additionally wanted to determine replication efficiency relative to the gene segment matched control, LS1D, in BC cells. AU-565, BT-474, ZR-75-1, and MCF7s were infected at an MOI of 0.1 with LS1D or S1/HER2nhd. Samples were collected at 0, 24, 48, 72, and 96 h p.i. and subjected to three freeze/thaw cycles. Plaque assays were performed on serial diluted lysates to determine virus titer at each timepoint. In each of the cell lines, S1/HER2nhd replication was not significantly different from LS1D, with both showing at least a three-log increase in viral titer by 72 h p.i. (Fig 7A-D). These findings suggest that modification of the σ1 protein via head domain replacement with HER2 peptide/Flag fusion does not significantly change the ability of the virus to replicate to high levels in BC cells. This indicates that this virus is likely to have the capacity to replicate and spread in BC tumors, expressing HER2 peptides at high levels in the infected cells, from lysis of infected cells, and from the surface of progeny virions produced and released during infection.

**Fig 7.**
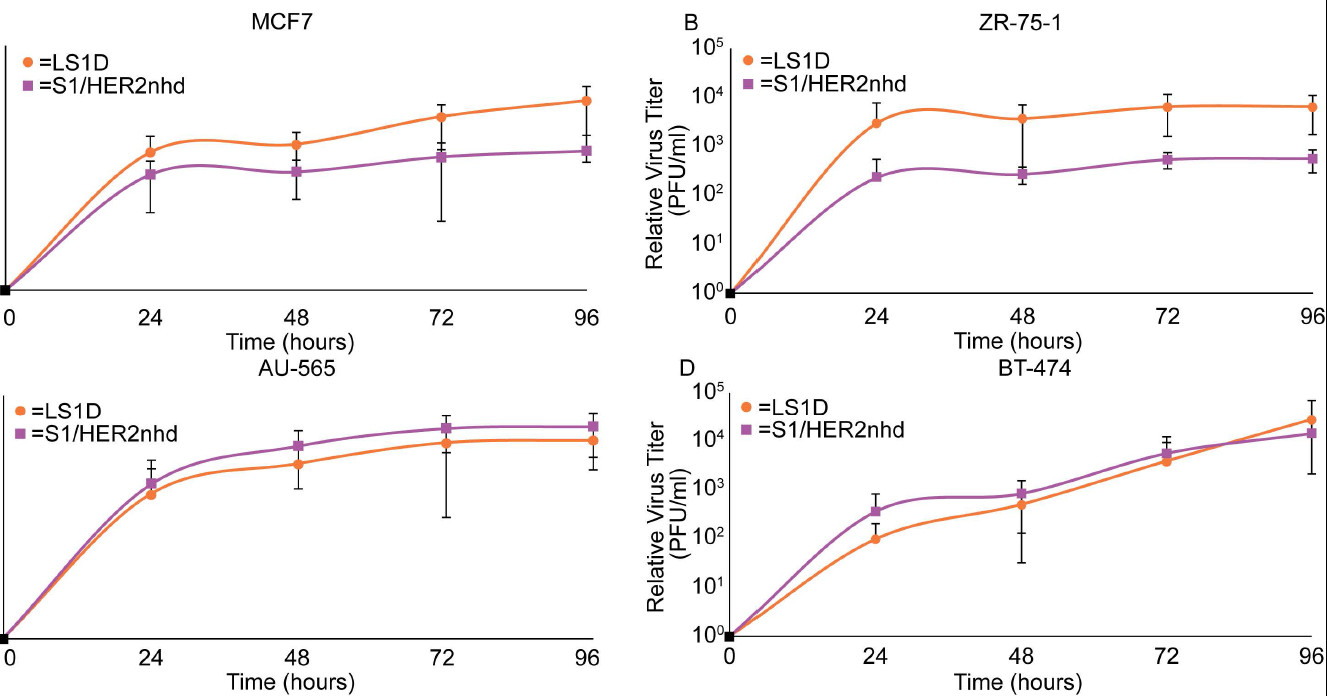
S1/HER2nhd and LS1D replicate to similar titers in BC cells. A) MCF7, B) ZR-75-1, AU-565, and D) BT-474 cells were infected with LS1D or S1/HER2nhd at an MOI of 0.1. At indicated time points, samples were subjected to three freeze/thaw cycles and plaque assays on L929 cells. The means and standard deviation from three experimental replicates are shown.

### S1/HER2nhd infection reduces viability in BC cells

The release of virus expressed HER2 peptide fusion proteins and progeny virus containing HER2 peptides on the virion particle is dependent at least in part on the ability of the virus to induce cell death and lysis. While our prior data show S1/HER2nhd replicates efficiently in BC cells, it does not specifically measure BC cell death induced during infection. To test this, each BC cell line was mock infected or infected with LS1D or S1/HER2nhd at an MOI of 1 for 24, 72, and 120 hours, at which time they were subjected to the CCK-8 assay which measures the reduction of tetrazolium salt via production of a formazin dye, WST-8, that is directly proportional to the number of live cells in the sample.

In MCF7 BC cells, which do not express high levels of HER2, both LS1D and S1/HER2nhd showed significant decreases in cell viability relative to mock infected cells by 24 hours, and viability further decreased at 72 and 120 h p.i (Fig 8A). In ZR-75-1 cells, which express moderate levels of HER2, both LS1D and S1/HER2nhd induced a significant decrease in cell viability by 120 h p.i. with LS1D inducing cell viability significantly more than S1/HER2nhd at this time point (Fig 8B). In the 2 BC cells that express high levels of HER2, AU-565 and BT-474, cell viability was significantly decreased by 120 h p.i. with LS1D and S1/HER2nhd infection relative to mock samples (Fig 8C, D). These results suggest S1/HER2nhd is able to induce cell death in BC cells with different genotypes, including those that overexpress HER2, at levels that are similar to a gene-matched control MRV.

**Fig 8.**
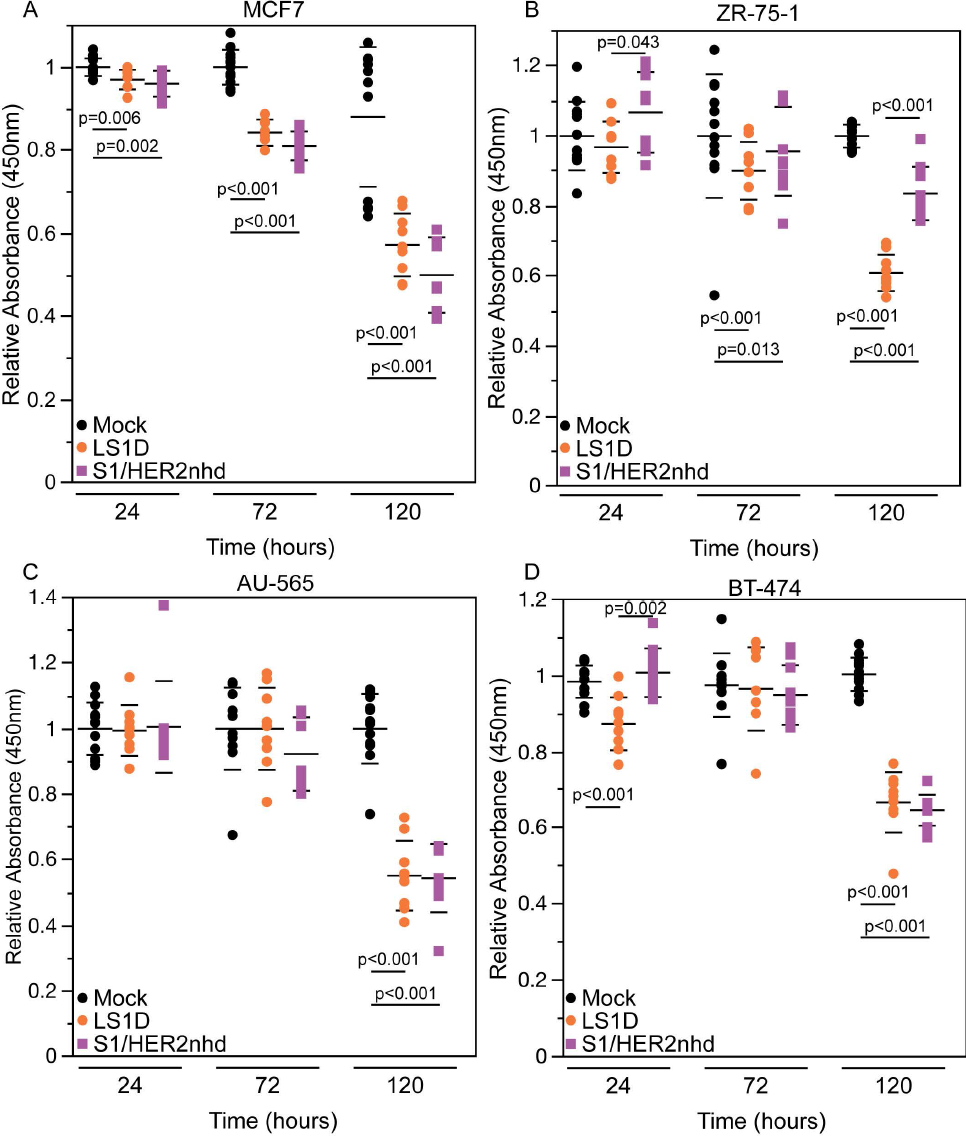
S1/HER2nhd reduces cell survival in BC cells. A) MCF7, B) ZR-75-1, C) AU-565, and BT-474 cells were mock infected or infected at an MOI of 1 with LS1D or S1/HER2nhd. At the indicated time points CCK-8 reagent was added to manufactures specifications and analyzed. All values were made relative to mock infected average at that timepoint. The means and standard deviation of three replicates are shown and significant differences between samples are indicated. P-values calculated using 2-sided student T test are shown.

## Discussion

In this work, we have constructed and rescued a novel recombinant MRV, S1/HER2nhd, and provided evidence that this virus expresses HER2 peptides as part of a σ1 fusion protein on the virion surface and inside infected cells. We have additionally shown that S1/HER2nhd replicates in and reduces cell viability in BC cells at levels similar to a WT gene-matched control virus. Using MRV, which has been shown in its WT form to stimulate an anti-tumor response, to deliver tumor associated antigens like HER2 directly to tumor cells is a natural progression in the search for ways to improve the therapeutic potential of the virus. This work provides additional support to published findings that creating recombinant viruses in the MRV backbone that are armed to improve oncolytic, immunotherapeutic, or subtype specific tumor targeting is feasible. However, much additional research is needed to bring this platform to the clinics. Of utmost importance will be investigating the extent that S1/HER2nhd is able to enhance a HER2 specific immune response and inhibit tumor growth in vivo in HER2+ BC animal models relative to WT MRV.

The work presented here, as well as in other studies, suggests that one critical hurdle that needs to be overcome when using MRV as a delivery vehicle for non-viral sequence is one of genetic stability. S1/HER2nhd, while stable for 3-6 passages, accumulates mutations or deletions upon further passage in vitro. Interestingly, each of the mutations/deletions in this work resulted in the loss of expression of the HER2 peptide/Flag tag fusion and maintained expression of the truncated σ1 protein. This suggests that the presence of the fusion sequence was selected against and may have been detrimental to virus growth. This pressure to mutate did not always involve loss of sequence at the genomic level, further suggesting that it was likely the fusion protein itself that was inhibitory. Previous studies have shown that a human GM-CSF gene added to an S1 head domain deletion mutant similar to that used in our study is also not stable (Kemp et al., 2019). Interestingly this group created a codon optimized human GM-CSF recombinant MRV, which also was not fully stable. However, a recombinant constructed in a nearly identical manner using murine GM-CSF was stable (Kemp et al., 2019). In these studies, the GM-CSF proteins were expressed using the 2A approach, which would result in 2 proteins, truncated σ1 and GM-CSF, from a single RNA. The 2A mechanism is not 100% efficient, and the production of σ1/GM-CSF fusion proteins may have contributed to instability of these viruses, although it is also possible that changes within the S1 gene segment RNA itself drove the loss of the GM-CSF. Our studies suggest that there are likely protein sequences or structures that alter protein folding and introduce instability into the individual recombinant viruses. If this is true, one approach to creating a more stable MRV that expresses HER2 peptides or other proteins may be to add a non-structured linker sequence between the WT MRV sequence and the exogenous sequence. This may resolve folding issues and increase overall stability of the genome such that gene fusions in S1 would continue to be expressed over many passages. Certainly, it is unlikely that as it currently exists that S1/HER2nhd would remain stable for long periods under the additional pressure of survival in the context of the host immune response. Therefore, resolving genome stability issues will be necessary to move recombinant viruses similar to S1/HER2nhd forward.

There have been a number of studies utilizing viruses or virus-like particles to stimulate an anti-HER2 immune response. This includes viruses or virus-like particles that contain and deliver the HER2 gene for expression, and virus-like particles that present HER2 on their surface. In each of these studies it was shown that the delivery of HER2 in gene or protein form resulted in an increase in the immune response against HER2. For example, HER2/neu-MPtVLPs, murine polyomavirus viral like particles that contain the extracellular and transmembrane domains of HER2 within the VLP, have been shown to elicit primarily a CD8+ T cell response to HER2, and an antibody response only to VP2, one of the viral proteins of murine polyomavirus used to generate the VLP (Andreasson et al., 2009; Tegerstedt et al., 2005). However, HER2-VLP, a viral like particle with the HER2 extracellular domain fused at the N-terminus to SpyCatcher (the CnaB2 domain from the fibronectin-binding protein FbaB or Streptococcus pyogenous protein binding partner), which presents the HER2 extracellular domain on the surface of the VLP, produced antibodies against HER2 when injected into mice grafted with mammary carcinoma cells expressing HER2 (Palladini et al., 2018). CD8+ T cells were also generated against HER2 when mice were injected with HER2-VLP (Palladini et al., 2018). This prior data suggests that our approach to create a virus that expresses immunogenic HER2 peptides within cancer cells and on the virus surface, on a virus known to preferentially replicate in tumor over non-tumor cells is likely to elicit a HER2 immune response that is inclusive of a B and T cell response from the infected tumor cells. Based on previous studies with the peptide vaccines, E75, GP2, and AE36, we expect at least a CD8+ T cell response that targets HER2 expressing cells (Holmes et al., 2008; Mittendorf et al., 2006). Even though previous studies all used the peptides in addition to GM-CSF, we still expect generation of CD8+ T cells with S1/HER2nhd infection, since WT MRV infection has been shown to increase GM-CSF levels (Gujar et al., 2010; Parakrama et al., 2020). Moreover, with GP2 in the transmembrane domain and AE36 found in the intracellular domain, we expect cells expressing the truncated p95HER2, which resists current FDA approved therapies for HER2+ BC, may also be targeted by CD8+ T cells generated by S1/HER2nhd. Additionally, there is potential for antibodies to be made against HER2 after infection with S1/HER2nhd, as was seen with HER2-VLP when the full extracellular domain of HER2 was fused onto the VLP surface (Palladini et al., 2018). During infection with WT MRV, neutralizing antibodies are generated that can bind σ1, (Dietrich et al., 2017). Therefore, we expect that S1/HER2nhd is likely to produce HER2+ specific CD8+ T cells, and HER2 specific antibodies when tested in vivo.

Our findings suggest that S1/HER2nhd binds and replicates in BC cells in vitro. The pathway followed by WT MRV, from intravenous injection to the tumor microenvironment is not well understood. It has been shown that MRV is taken up by peripheral blood mononuclear cells (PBMC)s during intravenous injection into patients (Adair et al., 2012). Additional studies have shown that MRV can be taken up by DCs and T cells during intravenous injection in mice, resulting in delivery of replication competent MRV to tumor cells (Ilett et al., 2009). In clinical trials, 81% of patient tumor samples are positive for MRV after intravenous delivery (Moore et al., 2019). This percentage increases to 96% of patient tumor samples if patients with melanoma and skin biopsies are excluded, suggesting intravenous delivery of MRV is highly capable of reaching tumor cells (Moore et al., 2019). Prior studies using a σ1 truncation mutant that is unable to bind JAM-A, similar to that in S1/HER2nhd, suggest that JAM-A binding is not necessary for MRV infection and killing of a number of tumor cells in culture (Kemp et al., 2019). Our data with S1/HER2nhd would support that JAM-A binding is also not necessary for binding and replication in BC cells. Although these studies suggest that a recombinant virus such as S1/HER2nhd will not be hindered for localization to and replication in tumors in vivo, additional experimentation will have to be performed to determine if this virus will have the capacity to migrate to, infect, replicate in, and reduce tumor burden in a HER2+ BC animal model compared to WT MRV.

## Conclusion

Taken altogether, the data presented in this work builds upon growing evidence that, while we have much to learn about modifying MRV for targeted oncolytic therapy, the genome and viral proteins can be modified such that they express non-viral sequences. This may include diagnostic markers to identify tumors or immune activators to increase the overall anti-tumor immune response as has been previously demonstrated, or specific tumor associated antigens, or antigenic peptides as we have demonstrated here. The majority of MRV recombinants created thus far appear to be able to replicate in and kill tumor cells, although many of them lack genomic stability when passaged. Additional work further defining how to successfully modify the MRV genome and proteins such that they better accommodate exogenous sequences will aid in the engineering of stable and effective MRV-based oncotherapeutics. The production of stable, recombinant oncolytic MRVs is expected to lead to an enhanced oncolysis generally, or against specific tumor subtypes, resulting in improved therapeutic options and outcomes for cancer patients.

## Acknowledgements

We would like to acknowledge fellow members of the Miller laboratory for their input and discussion. This work was supported by the following funding sources: NIH R15 CA202984, the Margaret B. Barry Cancer Research Award, and the Lloyd Cancer Award.

